# Precise optical probing of perceptual detection

**DOI:** 10.1101/456764

**Authors:** Gilad M. Lerman, Jonathan V. Gill, Dmitry Rinberg, Shy Shoham

## Abstract

Establishing causal links between patterns of neuronal activity and perception is crucial for understanding brain function. Electrical and optogenetic stimulation experiments demonstrated that animals can detect activation of a few neurons. However, these methodologies offer very limited control of ensemble activity and yielded highly divergent thresholds. Here, we use holographic two-photon (2P) optogenetic stimulation to probe the detection of evoked neuronal activity at cellular and single action potential resolution, with millisecond precision. We find that mice can detect single action potentials evoked synchronously across <20 olfactory bulb neurons, while ruling out detection of indirect effects using a novel optical sham-photostimulation technique. Our results demonstrate that mice are acutely attuned to sparse, synchronous ensemble activity signals, introducing order-of-magnitude revisions to earlier estimates of perceptual thresholds.

---

Sensory information is sent to higher brain areas from the mouse olfactory bulb’s output layers, whose ~50,000 projecting mitral and tufted cells (MTCs), spontaneously generate 10^5^-10^6^ action potentials (or ‘spikes’) per second. Yet, animals can detect minute vital cues within this strong background. What is the neural basis of this capacity and what are the minimal features of neural activity that can convey sensory information [1-5]? Odor-evoked responses in MTCs have a temporal precision of ~10 ms [6-8], and a powerful class of ideas for ‘noisy’ neural information transfer involves precise spatiotemporal synchronization or coordination of multi-neuronal ensemble patterns[9, 10]. However, the readout requirements for such coordinated population ‘codewords’ remain entirely unknown. Optogenetic techniques play a key role in causally linking specific genetically defined sets of neurons to perception and behavior [11], but current applications have been limited to spatially coarse manipulations. Precisely stimulating pre-selected distributed sets of neurons is currently only possible with holographic 2P optogenetics [12-20], a recent variant whose use remains limited and which has not yet been applied for probing the perceptual readout of neural ensembles. Earlier realizations [12-14, 17, 21-26] had an inadequately low temporal precision (10s – 100s of ms), and low excitation efficiency also limited excitation more superficially than the ~250-300 µm deep mitral cell layer.

## All optical imaging and photostimulation

We simultaneously performed 2P holographic optogenetic stimulation and 2P calcium imaging [12, 19] in the olfactory bulb of awake, head-fixed mice outfitted with a cranial window. To boost excitation efficiency and temporal precision, our system used a low-repetition rate, amplified laser source (at 1028 nm) for stimulation [15, 16, 18, 27], which was combined with a standard femtosecond oscillator at 920 nm for imaging (Fig. 1A). For a given average power *P_avg_*, an amplified laser source with low repetition rates, *f*, (100’s of kHz vs. ~80MHz in standard Ti:Sapphire oscillator) has a much higher peak power, and 2P excitation efficiency is inversely proportional to both the laser’s *f*, and pulse width, *τ*, as 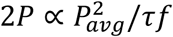 (equation 1 [28], Fig. S1). We used a spatial light modulator (SLM) to holographically generate soma-covering light patches (12-20 μm diameter, Fig. S2), targeted to excitatory, projecting MTCs and inhibitory granule cells (GCs) expressing ChrimsonR-tdTomato [29] together with GCaMP6s [30] (Fig. 1, B and C). This opsin-indicator combination resulted in densely co-expressing populations, enabling simultaneous optical readout and stimulation (Fig. S3).

**Figure 1.**
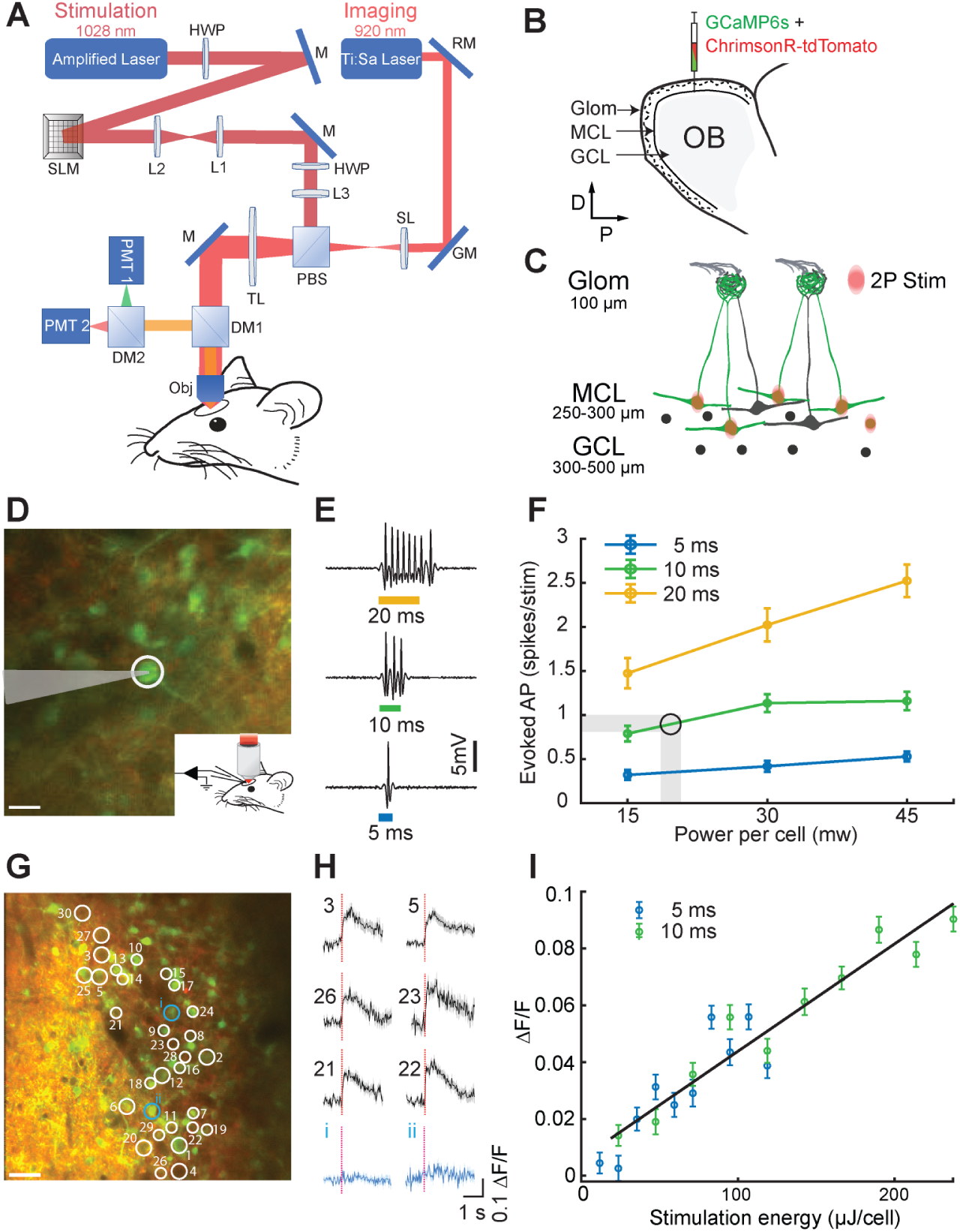
All optical imaging and photostimulation. **A.** Layout of the experimental system. The 2P imaging path is combined with the scanless holographic 2P photostimulation path for behavioral experiments in a head-fixed mouse with a cranial window. RM, resonant mirror; GM, galvanometer; SL, scan lens; PBS, polarizing beamsplitter; TL, tube lens; DM1, DM2, dichroic mirrors; PMT1, PMT2, photomultiplier tubes; HWP, half-wave plate; SLM, spatial light modulator; L1-L3, lenses; M, mirror; Obj, 16x objective. **B.** Injection of viruses into the mitral and the granule cell layers (MCL, GCL) in the olfactory bulb (OB). **C.** Schematic of 2P photostimulation patches targeted to the MCL of the OB. Large neurons, mitral cells; circles, granule cells; green, GCaMP6s response to photostimulation. **D.** 2P guided cell-attached electrophysiological recording in an awake mouse MCL coexpressing ChrimsonR-tdTomato (red) and calcium indicator GCaMP6s (green). The cell (white circle) was targeted by a light patch (scale bar, 20 µm). **E.** Examples of electrophysiological recordings during 5, 10 and 20 ms photostimulation, 628 kHz repetition rate. **F.** Average number of evoked spikes as a function of power per cell for different stimulation durations (mean ± s.e.m., *n*=5 cells). Shaded area and circle indicate the working zone chosen to elicit ≤1 evoked action potential. **G.** Thirty neurons were targeted for simultaneous 2P photostimulation (white circles) and 2P imaging (scale bar, 40 µm). **H.** Average GCaMP6s calcium response to photostimulation (red dashed line – stimulation onset) for 30 targeted cells (black) and 2 non-targeted cells (blue) (mean ΔF/F ± s.e.m., 30 trials, 10 ms stimulation duration, 0.06 mW/µm^2^,18 mW/patch). **I.** Average evoked calcium response as a function of photostimulation energy for 5 and 10 ms durations (mean ΔF/F ± s.e.m., *n*=20 cells, 40 trials).

To characterize the spiking response elicited by 2P photostimulation, we performed 2P guided cell-attached electrophysiological recordings (Fig. 1, D and F). Photostimulation of 5, 10 and 20 ms duration and average power levels of 15, 30 and 45 mW/patch (0.05, 0.1 and 0.14 mW/µm2 at objective output) generated short spiking epochs during illumination and immediately after stimulation offset (Fig. 1E, Fig. S4A). We saw a linear dependence of evoked spike count on stimulus duration (5, 10 and 20 ms illumination generated 0.41±0.06, 1.13±0.10 and 2.02±0.19. spikes, respectively)(Fig. 1F). Changing the average power between 15, 30, and 45 mW at a fixed duration of 10 ms yielded 0.79±0.09, 1.13±0.10, 1.15±0.10 spikes, respectively. The average latency for 10 ms, 30 mW photostimulation was 5.0 ms with a jitter of 3.1 ms (n=5 cells, 119 photostimulations), and stayed below 6.9 ms latency and 4.6 ms jitter for all power-time combinations, confirming rapid and temporally precise optogenetic stimulation (Fig. S4, B and C). Interestingly, this is not the system’s performance limit: 1 ms stimulations can elicit robust, tightly timed spiking (Fig. S4F), which together with the SLM’s fast (<3 ms) switching time enables >200 Hz photostimulation of different groups of neurons; MTCs can be driven at their natural firing rate (8-25 Hz) using trains of pulses, and up to 100 Hz (Fig. S4G). We next verified that moving the photostimulation patch away from the cell’s soma, both in the x-y plane (n=4 cells, 75 photostimulations), and along the z axis (n=9 cells, 150 photostimulations) ceased spiking activity, confirming minimal off-target excitation (Fig. S4, D and E), which could be further improved with new targeting strategies [13, 15, 18, 31]. For the behavioral experiments described below we chose to work with 10 ms and 18-20 mW/patch, which robustly elicited ~1 extra spike on average over the high tonic baseline activity (Fig. 1F, conservatively set at 0.9±0.1 spikes per stimulation to offset the impact of fluctuations and the saturating curve).

Next, we tested population-level photostimulation by holographically directing light patches to 30 neurons coexpressing ChrimsonR-tdTomato and GCaMP6s while recording fluorescence changes (30 trials, n=3 mice) (Fig 1, G to I). The total average power directed to the brain was <600 mW (10 ms x 18-20 mW/patch) and was sufficient to excite all targeted neurons (Fig 1H). Most non-targeted neuronal signals were not significantly modulated (blue traces), although, generally, current generation population-level calcium imaging cannot definitively preclude off-target and network-related activations [12, 16, 18, 26]. Target responses increased with stimulation strength, similar to our single-neuron electrophysiological observations (Fig 1I). In neurons simultaneously photostimulated with 5 and 10 ms durations (20 neurons, 40 trials/energy level, 800 photostimulations total, n=1 mouse), responses were linearly dependent on stimulation duration and increased monotonically with overall energy. This indicates that we can simultaneously deliver light to many cells in the field of view (FOV) and titrate the number of spikes evoked by the population of neurons targeted for behavioral interrogation.

### Behavioral detection of cellular-level photostimulation

We next tested the sensitivity of mice to ensemble photostimulation. To address this, we trained mice to detect precise photostimulation of sets of neurons and derived a direct single-spike resolution estimate of the detection threshold for olfactory bulb activity (Fig. 2, A to F). First, water-deprived head-fixed mice were trained to detect broad 1-photon optical stimulation (105 µm optic fiber coupled to 473 nm laser) of their olfactory bulb in a go/no-go paradigm (Fig. 2, A and B). Respiration was monitored, allowing photostimulation delivery to be timed to a particular respiratory phase (20 ms after inhalation onset) in a closed-loop manner. Mice reported detection of the stimulus by licking for a water reward, and withheld licking on trials without stimulation to avoid a punishment (5 sec. time-out) (Fig. 2B). Over the course of several sessions, laser power and duration were reduced 2,500-fold, while maintaining high detection performance (>70% correct (hits + correct rejects/total trials), n = 3) (Fig. 2D, Fig. S5A). We then transitioned mice to detect 10 ms, ~single spike 2P stimulation of a specific subset of 30 neurons (Fig. 2, C and D). Targeted neurons were co-localized putative MTCs or GCs (~15 MTCs + ~15 GCs), selected and targeted based on 1) co-expression of the opsin and indicator, and 2) reliability of responses across blocks and sessions (Fig. S6). Mice learned to perform the task with high accuracy over several sessions (6.7 ± 2 sessions (mean ± s.d.), n=3) (Fig. 2D, Fig. S5B). There was little variability in cellular responses to photostimulation across days, ruling out changes in performance related to virus expression, sensitization, or other non-behavioral factors influencing the efficacy of responses (Fig. S6).

**Figure 2.**
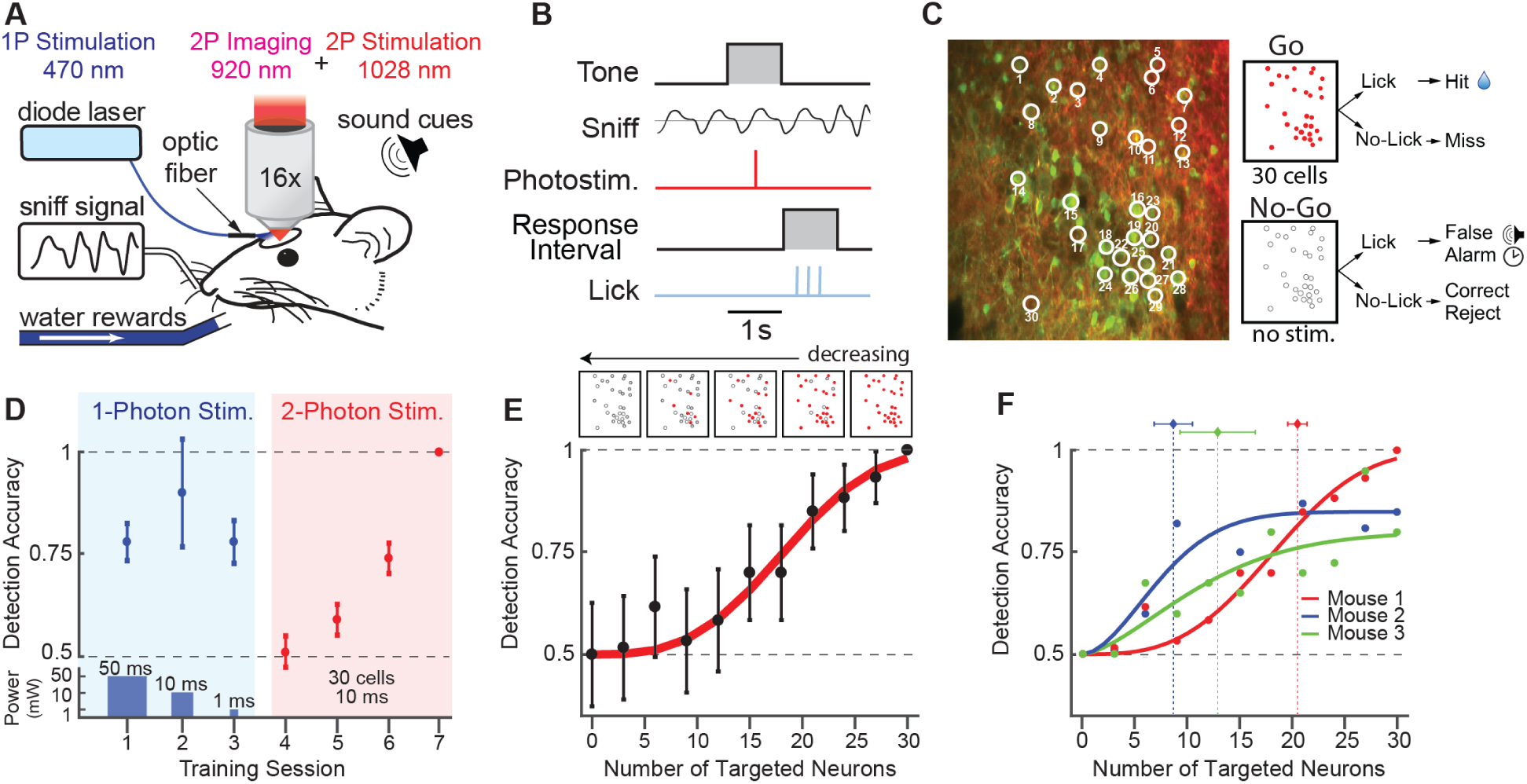
Behavioral detection of cellular-level photostimulation. **A**. Schematic of cellular photostimulation detection experiment. A head-fixed mouse with a chronically implanted window above the olfactory bulb is positioned in front of a lickspout and pressure sensor to monitor respiration (sniff). Neurons were stimulated with either 1-photon blue light (473 nm) generated by a diode laser, or the 2P holographic system. **B.** Trial structure for detection experiment. **C.** Left, neurons in the mitral cell layer (MTCs and GCs) coexpressing ChrimsonR-tdTomato (red) and GCaMP6s (green). Thirty neurons were targeted for simultaneous photostimulation (white circles). Right, outcomes for responses to the “go” and “no-go” stimuli. Red circles indicate stimulation of a particular cell, while empty circles indicate no stimulation. **D.** Detection performance in a representative training sequence. Mice were first trained to detect 1-photon photostimulation with decreasing energy from session to session (top, blue background) (mean ± 95% confidence intervals, *n*=1 mouse). Power and duration of 1-photon photostimulation (bottom, blue background). Then, mice were trained to detect 2P photostimulation of the same 30 targeted cells in each session (pink background) (mean ± 95% confidence intervals, *n*=1 mouse, 0.06 mW/µm^2^ intensity). **E.** Bottom, example detection accuracy for 2P photostimulation as a function of the number of targeted neurons, fit with a psychometric function (mean ± 95% confidence intervals, 60 trials per data point, *n*=1 mouse, 0.06 mW/µm^2^ intensity). Top, schematic of target stimuli used in experiment. **F.** Detection accuracy and threshold estimates across mice (60 trials/data point, mice 1 and 2, 30 trials/data point, mouse 3, *n*=3 mice, vertical dotted lines are Weibull threshold parameter fits ± bootstrapped 95% confidence intervals).

We further explored this sensitivity by testing detection performance while parametrically varying the number of targeted neurons. For stimuli, we generated holograms targeting subsets of neurons from the original 30 neuron pattern, while maintaining the same average power per neuron (0-30 cells targeted, 19±1 mW/cell). Detection performance varied monotonically as a function of the number of neurons targeted (Fig. 2E). Different mice exhibited different degrees of sensitivity, with threshold performance varying from 8.7±1.8 neurons at the minimum to 20±0.9 at the maximum (maximum likelihood estimate Weibull fits ± s.e.m. of bootstrap analysis) (Fig. 2F). Together, these data suggest that mice are exquisitely sensitive to changes in the synchronous activation of <20 olfactory bulb neurons, on the order of several spikes.

### Pulse duration tunability enables sham photostimulation control

To rule out behavioral detection of confounds, such as heat or indirect sensory stimulation (e.g. visual, tactile, etc.) we performed a control experiment using sham stimulation which reproduced all features of the stimulus but didn’t evoke spiking (Fig. 3, A to D). To achieve this, we increased the laser pulse duration from ~200 fs used for effective photostimulation, to >15 ps for sham photostimulation, thereby reducing the 2P signal by two orders of magnitude (equation (1)) yet maintaining the same average power, and thus all other aspects of the stimulus (Fig. 3, A and B; Fig. S7A). We tested the sham photostimulation ability to evoke spiking across a range of power levels and durations. Firing rate during illumination did not differ from baseline across targeted neurons for all power levels and durations (n=4, Wilcoxon rank sum test, P=0.99) (Fig. 3C, Fig. S7B) indicating that increasing the pulse duration abolished spike generation.

**Figure 3.**
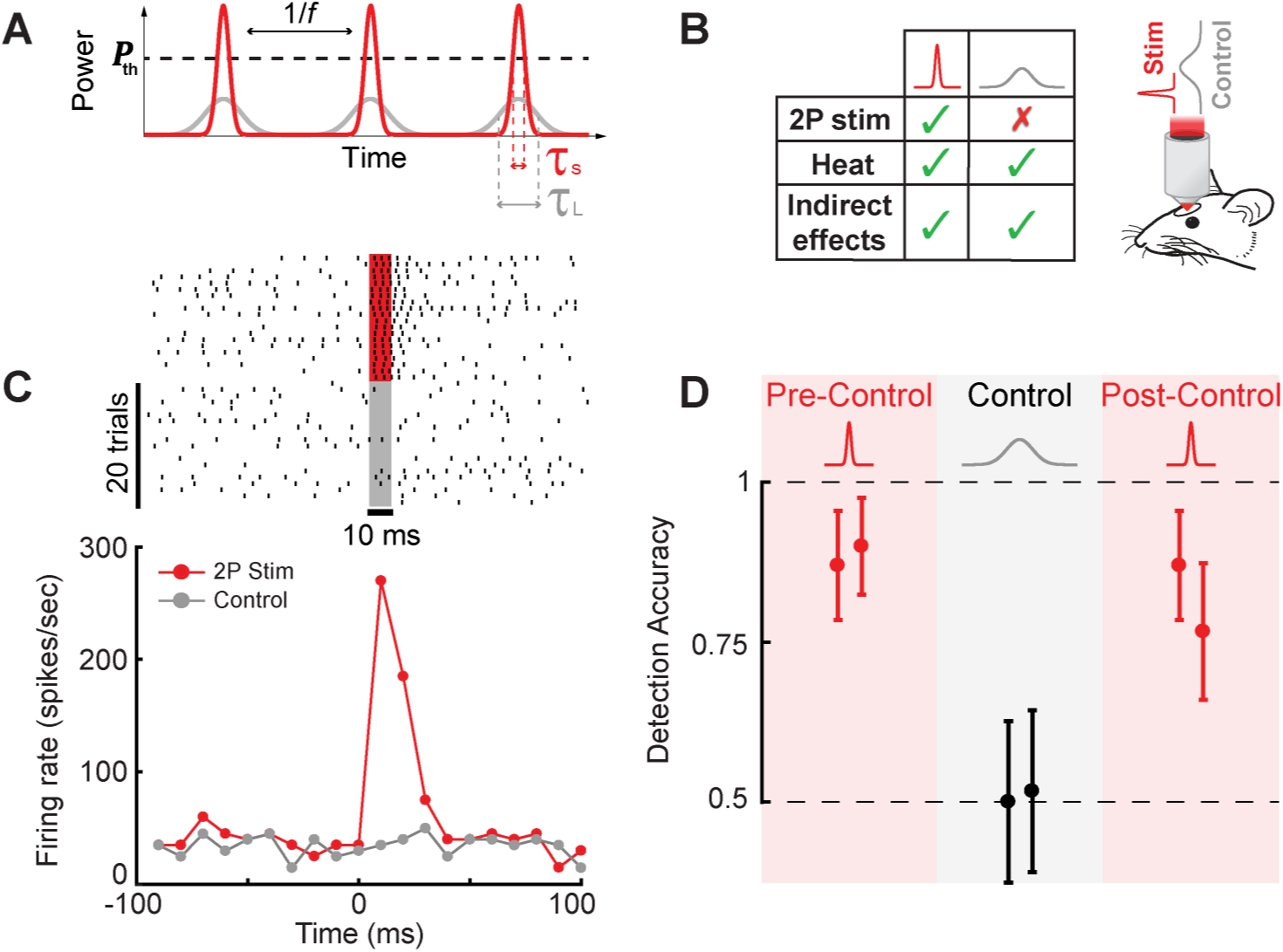
Pulse duration tunability enables sham photostimulation control. **A.** A schematic showing the effect of tuning the pulse duration. To photostimulate a cell, 2P excitation must exceed a certain threshold, ***P_th_***. As the pulse duration increases (*τ*_L_>*τ*_S_), the peak power, and therefore the 2P signal, decreases below the threshold and cannot stimulate the cell. **B**. Left, a table summarizing the differences in effects evoked by the short and long pulse duration stimuli. Right, a schematic of the behavioral setup for the sham photostimulation control experiment. **C**. Representative example raster plots (top) and PSTHs (bottom) for short pulse photostimulation and long pulse sham photostimulation (control) (red, short pulse photostimulation; gray, long pulse control; 20 trials per condition, n=1 cell, 30 mW, 10 ms illumination). **D**. Detection accuracy as a function of photostimulation condition. Blocks of sham photostimulation were interleaved between high-performance blocks. During the sham-stimulation blocks, detection accuracy dropped to chance level and was significantly different from both pre and post-control measurements (30 targeted cells, 0.06 mW/µm2, n = 2 mice, Fisher’s exact test, P<0.001).

Can mice detect the sham stimulation? We interleaved sham stimulation blocks among the highest-performance daily behavior sessions (Fig. 3D). Detection performance dropped to chance level, significantly below preceding and subsequent blocks (n=2, Fisher’s exact test, p<.001), demonstrating that photostimulation detection performance is determined by the spiking of targeted neurons and not confounding factors, despite the mouse’s motivation to utilize any informative cue.

## Discussion

Precise manipulation and monitoring of neural activity is critical for deciphering the behaviorally relevant features of the neural code. Previous experiments have lent support to the idea that small groups of sensory neurons may have a salient impact on perception [1-5], yet previous approaches left unclear the degree to which animals are attuned to the precise ensemble activity of sensory regions. This study provides the first direct evidence for the number of neurons and spikes necessary for photostimulation detection in the mouse, refining previous estimates by an order of magnitude (<30 vs. ≥300[2]). Remarkably, single spikes evoked synchronously across only a few neurons were detectable on top of the relatively high baseline firing rate of the olfactory bulb (8-25 Hz) [6-8], suggesting that mice might be extremely sensitive to not just the number, but the precise timing of behaviorally relevant neurons. In a 10ms time bin, olfactory MTCs spontaneously generate ~5000 spikes, and the ability to robustly and very quickly learn to detect <<1% ‘relevant’ spikes highlights a remarkable adaptive performance characteristic of sensory systems. Future work comparing thresholds derived using this technique across sensory systems, and cell type, will help further constrain models linking ensemble activity to sensory coding.

## Acknowledgments

The authors thank G. Serrano for surgical assistance, C. Wilson for aid in experimental programming, E. Chong for help with behavioral training, as well as A. Resulaj and H. Ma for help in behavioral system design. The authors also thank G. Kosche, M. Picardo, S. Benezra, and K. Martin, for assistance with juxtacellular recordings, as well as J. Little, S. Rosen, and D. Razanski for comments on the manuscript. **Funding:** This work was supported by the NIH BRAIN Initiative Grant (1U01NS090498-01), J. G. was supported by NIH graduate training grant (T90DA043219)

## Author contributions

D.R. and S.S. designed and initiated the study and supervised the project. J.G. and G.L. built the experimental system, carried out the experiments and performed the data analysis. All authors discussed the results and wrote the manuscript.

## Declaration of interests

The authors declare no competing interests

## Data and software availability

The datasets generated during and/or analyzed during the current study are available from the corresponding author on reasonable request.

## Methods

All procedures were approved under a New York University Langone Health institutional animal care and use committee (IACUC) protocol 161211-02. Male and female C57/Bl6 mice (*Mus musculus*, Jackson Laboratories) between 2 and 4 months old were used in all experiments and handled in accordance with institutional guidelines.

### Surgical preparation

Mice were anesthetized with isoflurane during viral injections and surgical implantation (2.0% during induction, 1.5% during surgery). A circular craniotomy was performed to expose both hemispheres of the dorsal olfactory bulb (3 mm craniotomy extending from the rostral rhinal vein to the naso-frontal suture, centered on the midline) using an air-driven dental drill (Midwest Tradition, FG 1/8 drill bit). Adeno-associated viral (AAV) vectors encoding the calcium indicator GCaMP6s (AAV5-Syn-GCaMP6s-WPRE-SV40, UPENN vector core) and red-shifted opsin ChrimsonR (AAV5-Syn-ChrimsonR-tdTomato, UPENN vector core) were mixed at a ratio of 1:1 and injected bilaterally using a stereotactic syringe pump (World Precision Instruments Inc.) at a rate of 0.1 µl min^-1^ (800 nl per hemisphere, 400-600 µm deep). Following injection, a cranial window was implanted replacing a circular piece of skull by a glass coverslip (3 mm diameter, Warner Instruments) that was secured in place using a mix of self-curing resin (Orthojet, Lang Dental) and cyanoacrylate glue (Krazy Glue). For electrophysiological recordings, the cranial window contained two ~1 mm diameter holes pre-drilled and filled with a silicone elastomer (Kwik-Sil, World Precision Instruments). A custom 3D-printed headpost [32] was placed around the cranial window and affixed to the skull using C&B Metabond dental cement (Parkell). Each animal recovered for at least 10 days prior to experiments.

### Imaging and photostimulation

The system combined an imaging arm and a photostimulation arm to achieve all-optical two-photon imaging and photostimulation. Imaging was performed on a custom multiphoton microscope system based on the MIMMS 1.0 design (HHMI Janelia Research Campus, Ashburn, VA). Two-photon fluorescence of GCaMP6s and tdTomato was excited at 920 nm using a mode locked, femtosecond-pulsed, Ti:Sapphire laser with dispersion compensation (Mai Tai eHP DS, Spectra-Physics, Mountain View, CA). The beam was relayed and magnified by a telescope (scan lens, f=35 mm, and tube lens, f=200 mm) to the back-aperture of a 16×/0.8-NA water immersion objective lens (Nikon) mounted to a piezoelectric stage (P-725KHDS, Physik Instrument). Images were acquired at 30 Hz using resonant-galvanometer raster scanning (Cambridge Research). Emitted photons were reflected by a dichroic mirror (DM1, FF705-Di01, Semrock) and separated to either green or red channels via a dichroic mirror (DM2, 565dcxr, Chroma) and fluorescence was detected using GaAsP photomultiplier tubes (H10770PB-40, Hamamatsu). Images were digitized and recorded using ScanImage software [33] (Vidrio Technologies). The stability of the imaging and head-fixation system permitted imaging of the same FOV over multiple weeks in longitudinal experiments.

Two-photon photostimulation was performed using an amplified femtosecond laser source (Pharos, SP-06-600-PP Light Conversion, 6W maximal average power, 190 fs pulse width at 628 KHz repetition rate, except where noted) at 1028 nm. The repetition rate and pulse duration of the laser were tunable and were used to find the optimal parameters for 2P excitation, and to change the laser’s peak power for the control experiments. The beam power was controlled via the laser’s software and was measured at the laser output and also after the objective lens to verify the power level hitting the brain. We controlled the internal Pockels cell of the laser with a TTL pulse and used it as the shutter for the system achieving very short <1ms opening times not possible with a mechanical shutter. The beam was expanded with a telescope (BE02M-B, Thorlabs) to fill the SLM active area and the polarization was optimized with a half-wave plate (HWP, AHWP10M-980, Thorlabs) to achieve the best efficiency of the SLM. The reflective SLM (HSPDM512–920-1110, Meadowlark Optics, optimized for 1064 nm, 7.68*7.68 mm active area, 512x512 pixels), has an overdrive mode with a 3 ms refresh time, so combined with the rapid photostimulation approach outlined in this paper, photostimulation using different holographic patterns could be achieved in less than 10 ms. After acquiring the desired phase profile, the beam was relayed by a 1:1 telescope (L1, f=100 mm, L2, f=100 mm) to a mirror. Blocking of the zeroth diffraction order was done at the focal plane of lens L1 using a piece of aluminum foil that was glued onto a glass coverslip. An additional telescope with a ~ 1:3 ratio (L3, f=75 mm, and the tube lens, f=200 mm) was used to magnify and project the beam onto the objective lens aperture. The imaging and photostimulation paths were combined by a polarizing beamsplitter cube (PBS253, 900-1300 nm, Thorlabs) and an additional HWP (AHWP10M-980, Thorlabs) was placed before the beamsplitter to rotate the polarization of the diffracted beamlets to maximize power going through the polarizing beamsplitter.

Phase masks were loaded to the SLM using the manufacturer’s software development kit (SDK) and custom experimental control software written in Python. This allowed us to select from a set of pre-generated phase masks depending on the conditions of the experiment, permitting presentation of randomized and counterbalanced stimuli. Phase calculations were performed using custom MATLAB software that made use of a modified Gerchberg-Saxton (GS) algorithm [34] to optimize the intensity distribution at the focal plane. We mapped efficiency across the photostimulation FOV by generating phase masks to translate a single light patch across the FOV. We measured the intensity at the objective output for each hologram and calculated the relative efficiency of the holograms as a function of position. This characterization was later used to compensate for efficiency variations as we weighted the desired intensity pattern with this efficiency map. The registration of the photostimulation FOV and the two-photon imaging FOV was performed before each experimental session to ensure precise cellular targeting. A calibration pattern was burned onto a fluorescent plate (Ted Pella, INC.) by the photostimulation system. The plate was imaged by the 2P imaging system and the calibration pattern was used for the registration of the two systems.

The point spread function (PSF) of the light patches was characterized using a widefield microscope. A thin (<5 μm) fluorescein layer was illuminated with a light patch, and the excited fluorescence signal was collected through the objective lens, the tube lens and a mirror (MGP01-350-700, Semrock) onto a CMOS camera (DCC1240M – GL, Thorlabs). The axial dimension of the photostimulation PSF was measured by generating holograms of the same light patch in different axial positions spanning the focal plane and measuring the excited fluorescence signal for each hologram. This technique was also used to find the effective axial resolution of the photostimulation system by measuring the GCaMP6s calcium response of targeted neurons to each axial value. This measurement used a considerably higher power per patch than was used for behavioral experiments, 30 mW/patch instead of ~19 mW/patch for behavioral detection, in order to boost signal-to-noise for optical measurements, though this is likely to have increased the estimated full width at half max for the effective axial PSF. This widefield path was also used for coarse alignment of the imaging system around the same FOV in the behavioral experiments.

### Online motion correction

For behavioral experiments that required targeting the same neurons for photostimulation across days, we implemented an online motion correction method. In these experiments, the FOV was first aligned to a reference image manually, then the position was fine-tuned automatically using a closed-loop algorithm. This algorithm attempted to minimize the difference between the reference image and the FOV by iteratively moving the microscope stage (Sutter 285) to reduce the residual displacement computed using a rigid motion correction package (NoRMCorre, Flatiron Institute [35]). In addition to aligning the FOV across days, we performed this routine between blocks during each behavioral session to minimize the effect of slow x-y drift due to brain and microscope motion, therefore ensuring the photostimulation targets remained consistent throughout each session.

### Image processing and data analysis

Data analysis was performed using custom-written software in ImageJ (NIH) and MATLAB (MathWorks). Images were aligned to a session-averaged template image using a non-rigid motion correction package (NoRMCorre). Cellular regions of interest (ROIs) were manually drawn using both GCaMP6s and ChrimsonR-tdTomato channels, and raw fluorescence time-courses were extracted. For each cellular ROI, we subtracted the mean fluorescence of the surrounding neuropil to reduce the influence of any contamination and computed Δ*F*/*F*0. We classified neurons as responsive if they showed a significant difference in their average fluorescence signal immediately after photostimulation versus spontaneous activity preceding photostimulation (*P* < 0.05, Two-sample Kolmogorov-Smirnov test).

### Targeted juxtacelluar electrophysiology

*In vivo* juxtacellular recordings were performed during 2-photon imaging for guided access to olfactory bulb neurons. Recordings were performed with a Multiclamp 700b amplifier (Molecular Devices) and digitized at 20 kHz using an FPGA (National Instruments) and Wavesurfer software (Janelia Research Campus). Borosilicate glass pipettes were pulled to 5-11 MΩ (~1-1.5 µm tip size) and filled with external solution (130 mM K-gluconate) containing fluorescent dye for visualization (Alexa 488/594 mixture, Milipore). Before recordings, silicone plugs were removed from the cranial window and the dura was continuously perfused with sterile saline solution. After each recording session, holes in the cranial window were re-sealed using silicone elastomer, permitting multiple recording sessions for each animal.

### Behavioral training

All behavioral events (imaging time-stamping, respiration monitoring, stimulus delivery, water delivery, and lick detection) were monitored and controlled by custom programs written in Python interfacing with a custom behavioral control system (Janelia Research Campus) based on an Arduino Mega 2560 microcontroller. Photostimulation detection training began after at least 7 days of water restriction (1ml/day), and at least 14 days after viral injection. Mice were housed on a reverse light/dark cycle, and training took place during the day. Training began with habituation to head-fixation (1-2 days), while animals were free to run on a 3D printed wheel. This was followed by water-sampling sessions (1-3 days) in which mice learned to lick a tube to receive 2 µl droplets, until they learned to lick enough to receive the full 1ml of water for the day. Licking was detected using a capacitive touch sensor (Sparkfun) coupled to hypodermic tubing which triggered the release of water droplets by a pinch valve.

Once the animals reliably licked, they were placed on a photostimulation detection task to shape their behavior, in which mice reported the presence or absence of photostimulation presented broadly to one hemisphere of the olfactory bulb. Photostimulation was delivered using a continuous wave diode laser (FTEC2, BLUE Sky Research) at 473 nm through a 105 µm core diameter optic fiber (N.A. 0.22, beam diameter ~1.0mm) placed over the cranial window at a 53° angle. Animals performed a go/no-go task, in which ‘go’ (photostimulation) and ‘no-go’ (no photostimulation) trials were pseudo-randomly interleaved in a block structure (60 trials/block, 30 ‘go’, ~600 trials/day). Each trial consisted of a pre-stimulus period (500 ms), stimulus period (20-70 ms), a delay (500 ms), response period (500 ms), and an inter-trial interval (ITI) with variable duration (5-7 s). On all trials, a broadband tone (piezo buzzer) and masking LEDs signaled the start of the pre-stimulus period and persisted through the stimulus period and delay. The masking LEDs were chosen such that the central wavelength matched that of the diode laser (473 nm) and were positioned near the animals’ eyes to prevent detection based on visual cues. Offset of the tone and masking LEDs signaled the response period during which an animal could lick to receive a reward on “go” trials or withhold licking to avoid a punishment on “no-go” trials. Incorrect licking (false alarms) during the response period resulted in a tone from a second piezo buzzer of a different frequency (500ms) and a 5s “time-out” added to the ITI. During all other epochs animals were neither punished nor rewarded for licking, and after initial training mice generally did not lick outside of the response period.

Respiration was monitored using a pressure transducer coupled to a custom Teflon “odor port”, which continuously passed filtered air over the mouse’s nostrils at a rate of 1 l/min. Photostimulation was delivered at a fixed delay (20 ms) after the onset of inhalation during the stimulus period on “go” trials. This timing corresponded roughly to the onset of the earliest odor driven responses reported by previous experiments[7], so this period was used to avoid interference from potentially uncontrolled olfactory stimulation. Further, previous experiments targeting olfactory circuits have revealed that mice are highly sensitive to the respiration phase at which optogenetic photostimulation is presented [36, 37]. Therefore, we fixed this parameter to avoid any confounding effects timing might contribute to behavioral detection.

During shaping, mice were initially trained to detect relatively strong photostimulation (40 mW, 50 ms duration, 473 nm). Once performance for a given pulse energy exceeded 0.75 proportion correct, the pulse energy was decreased iteratively until mice performed above 0.75 proportion correct at 1 mW for 2 ms duration for at least one block. The fiber was positioned above the same hemisphere of the olfactory bulb across days, though not in the exact same position. In subsequent sessions, mice transitioned to detecting 2-photon photostimulation of 30 neurons within the trained hemisphere (18-20 mW/cell, 10 ms duration). All experimental timing and cues were identical to the shaping task, except that the masking LED was no longer used. The 30 neurons were held constant across days and confirmed to be responsive to photostimulation outside of the behavioral task in a block prior to each training session (30 repetitions, 3-5 s I.S.I.).

Once performance on the 30-cell detection task plateaued, animals were tested on detection performance as a function of the number of targeted neurons. In this experiment 10 holograms were generated by progressively removing targets from the trained pattern. Removed targets were chosen at random without replacement. Detection performance was tested by pseudo-randomly interleaving blocks of each reduced hologram.

### Optical sham-stimulation control

To confirm that detection relied on the spiking of neurons, and not confounding factors, animals were also tested on a control “sham stimulation” condition. The control stimulus was achieved by increasing the photostimulation pulse width from ~200 fs, to >15 ps by adjusting the compressor. The laser’s average power (and therefore the delivered heat and indirect sensory effects that might be evoked) was held fixed. Mice experienced blocks of the sham stimulation interleaved with blocks of normal stimulation during the go/no-go stimulation detection task after the task had been learned (>75% detection accuracy).

### Behavioral analysis

Results of behavior were analyzed in custom-written MATLAB programs with behavioral performance quantified in terms of proportion of correct trials. To determine each mouse’s sensitivity to the number of targeted neurons, psychometric performance curves were fit using a Weibull function and maximum likelihood estimation. Error bars for parameter estimates were generated using a bootstrap analysis.

### Statistical analysis

All statistical analyses were performed using MATLAB. The effect of sham-photostimulation on spiking activity was determined using a Wilcoxon rank sum test (Fig. S7B). To compare behavioral performance during normal photostimulation and sham-stimulation conditions, statistical significance was assessed using Fisher’s exact test (two-sided) (Fig. 3D).

## Supplemental Figures

**Figure S1.**
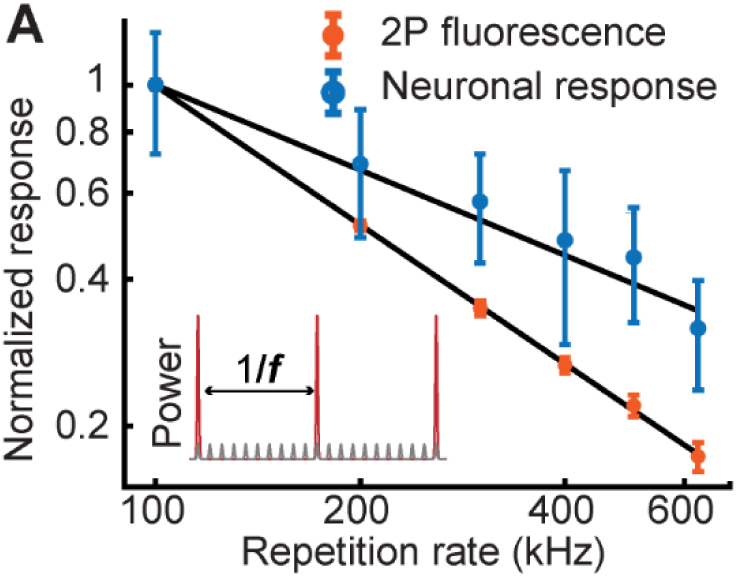
Characterization of 2P excitation dependence on repetition rate. **A.** Evoked 2P fluorescence signal measured using fluorescein, and neuronal GCaMP6s responses as a function of the laser’s repetition rate for a constant average power (mean ΔF/F ± s.e.m., *n*=10 cells, 30 stimulations per rep. rate). The data is normalized to the mean response value at the lowest repetition rate. Inset shows a schematic of the laser pulses for high and low repetition rates.

**Figure S2.**
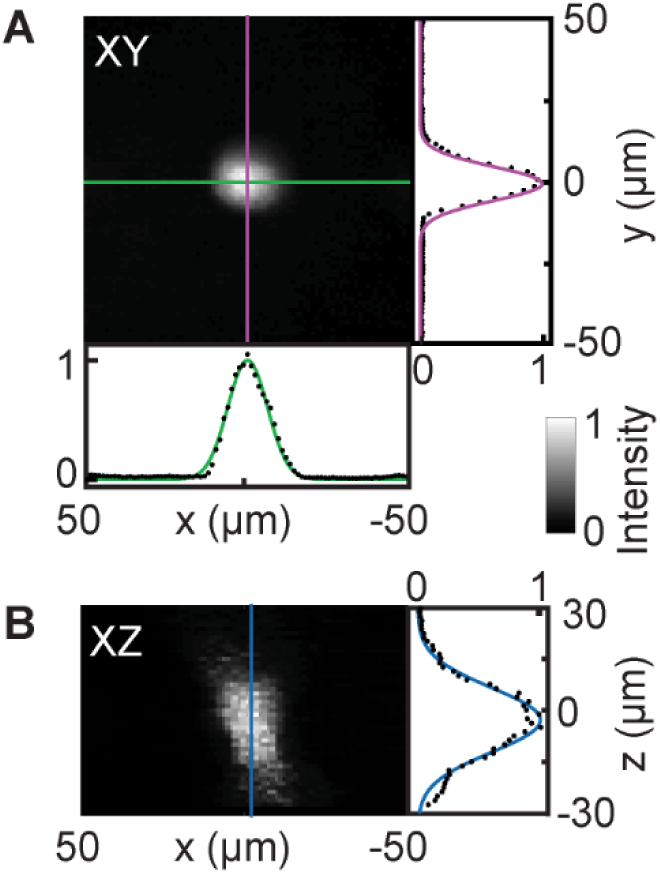
Photostimulation point spread function. Light patches generated using the Weighted Gerchberg-Saxton algorithm [34] were characterized using a widefield microscope. A thin (<5 μm) fluorescein layer was illuminated with a light patch and the 2P excited fluorescence signal was projected onto a CMOS camera. **A-B.** The photostimulation point spread function measured 15 μm and 12.5 μm full width at half max in the x and y dimensions and 23.7 μm in the z dimension.

**Figure S3.**
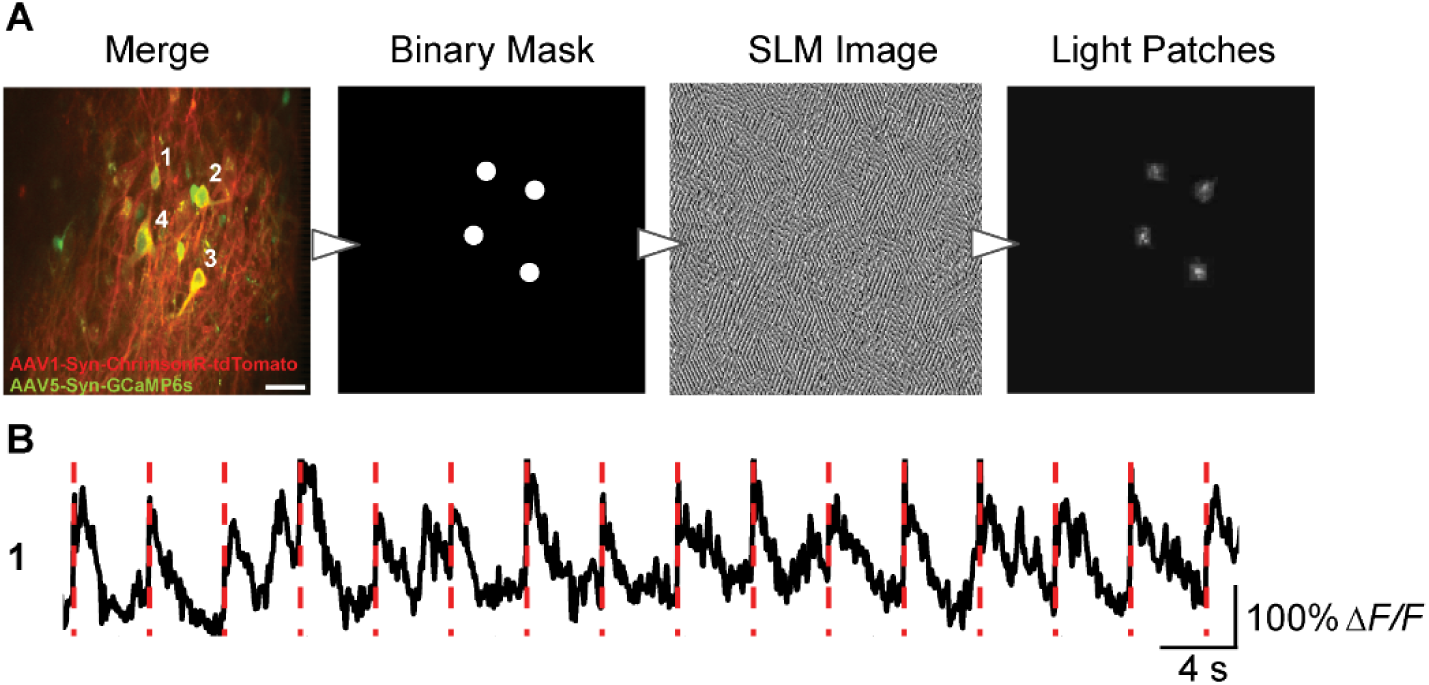
Holographically targeting multiple neurons with light patches. **A**. Illustration of workflow for holographically targeting light patches to multiple neurons. Scale bar, 30 µm**. B**. Example trial-by-trial calcium response to photostimulation for one neuron (4 targeted cells, 50 ms photostimulation, 16 trials, 0.1 mW/µm^2^).

**Figure S4.**
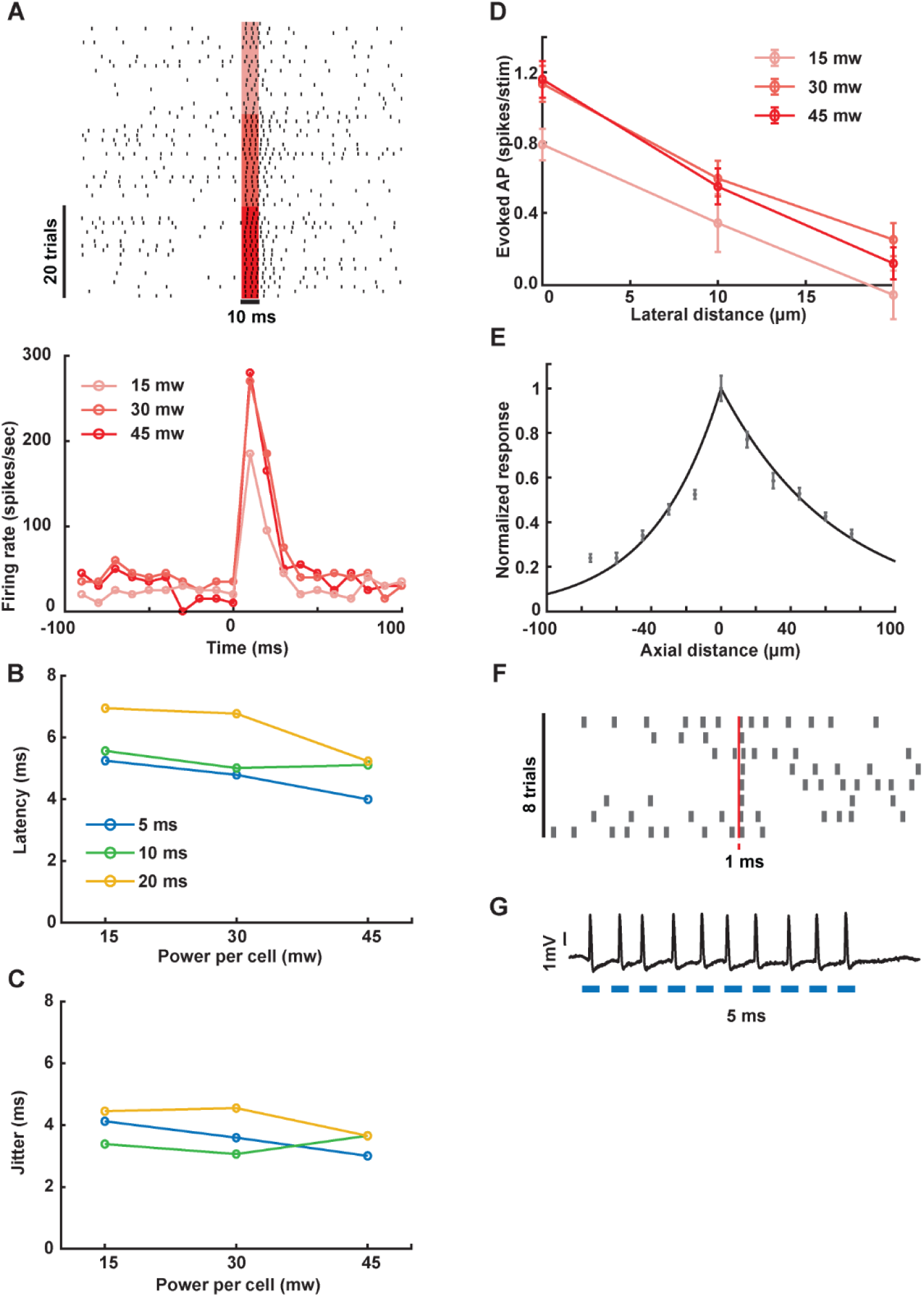
Electrophysiological characterization of photostimulation. **A.** Example raster plots (top) and correspondent PSTHs (bottom) for different stimulation powers, (red shading, photostimulation on; *n*=1 cell, 20 trials per condition). **B-C.** The latency (**B**) and jitter (**C**) of evoked spikes as a function of power per cell for different stimulation durations. The latency is defined as the time from photostimulation onset to the first spike within a 20 ms window, and jitter is defined as the latency’s standard deviation (*n*=5 cells). **D-E.** The efficiency of off-target photostimulation when moving the light patch away from the cell’s soma laterally (**D,** mean ± s.e.m., n=4 cells) and along the z-axis (**E,** mean ± s.e.m., n=9 cells, 0.1 mW/µm^2^). **F.** Raster plot of 1 ms photostimulation (45 mW, 0.14 mW/µm^2^, *n*=1 cell). **G.** 100 Hz photostimulation train (5 ms on / 5 ms off duty cycle, blue lines; 45 mW, 0.14 mW/µm^2^, *n*=1 cell).

**Figure S5.**
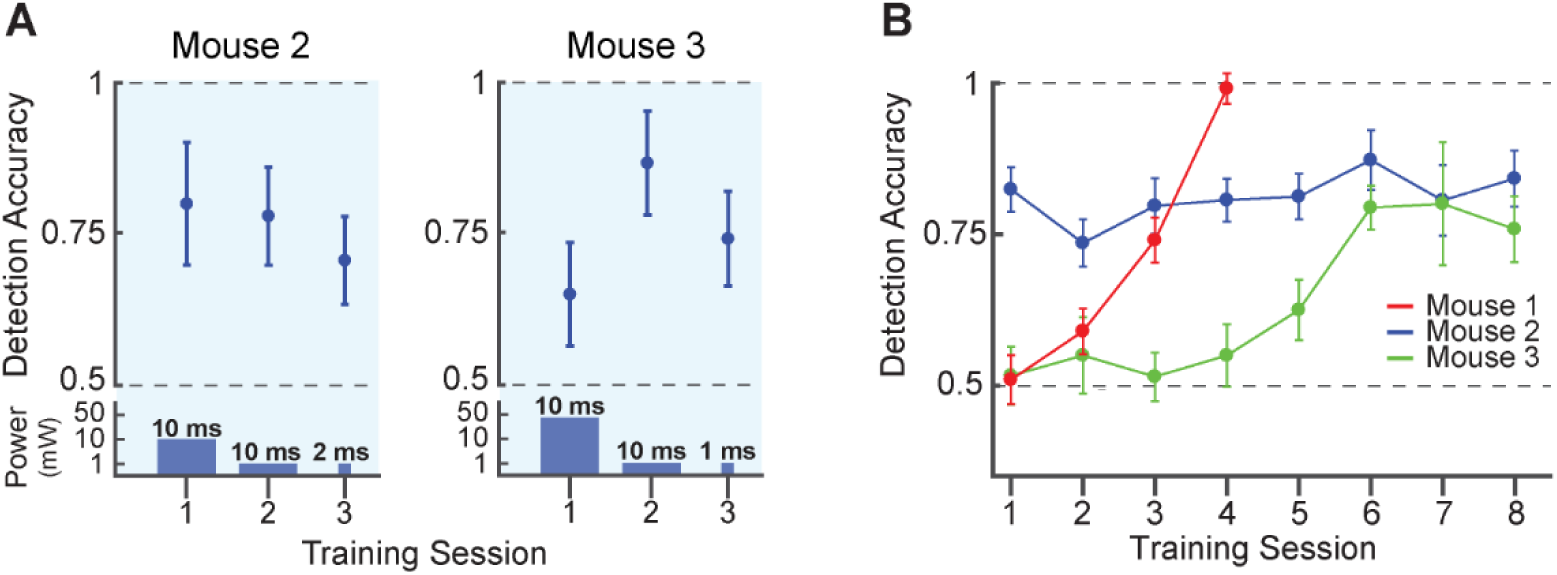
Photostimulation detection training. **A**. Performance for 1-photon photostimulation detection training for individual mice. 1-photon photostimulation energy was decreased from session to session. Power and duration of 1-photon stimulation is schematized at the bottom of the graph. **B**. Performance for 2P photostimulation detection of the same 30 cells across sessions (mean ± 95% confidence intervals, *n*=3 mice, 0.06 mW/µm^2^). All mice reached a performance >75% correct within 7 sessions.

**Figure S6.**
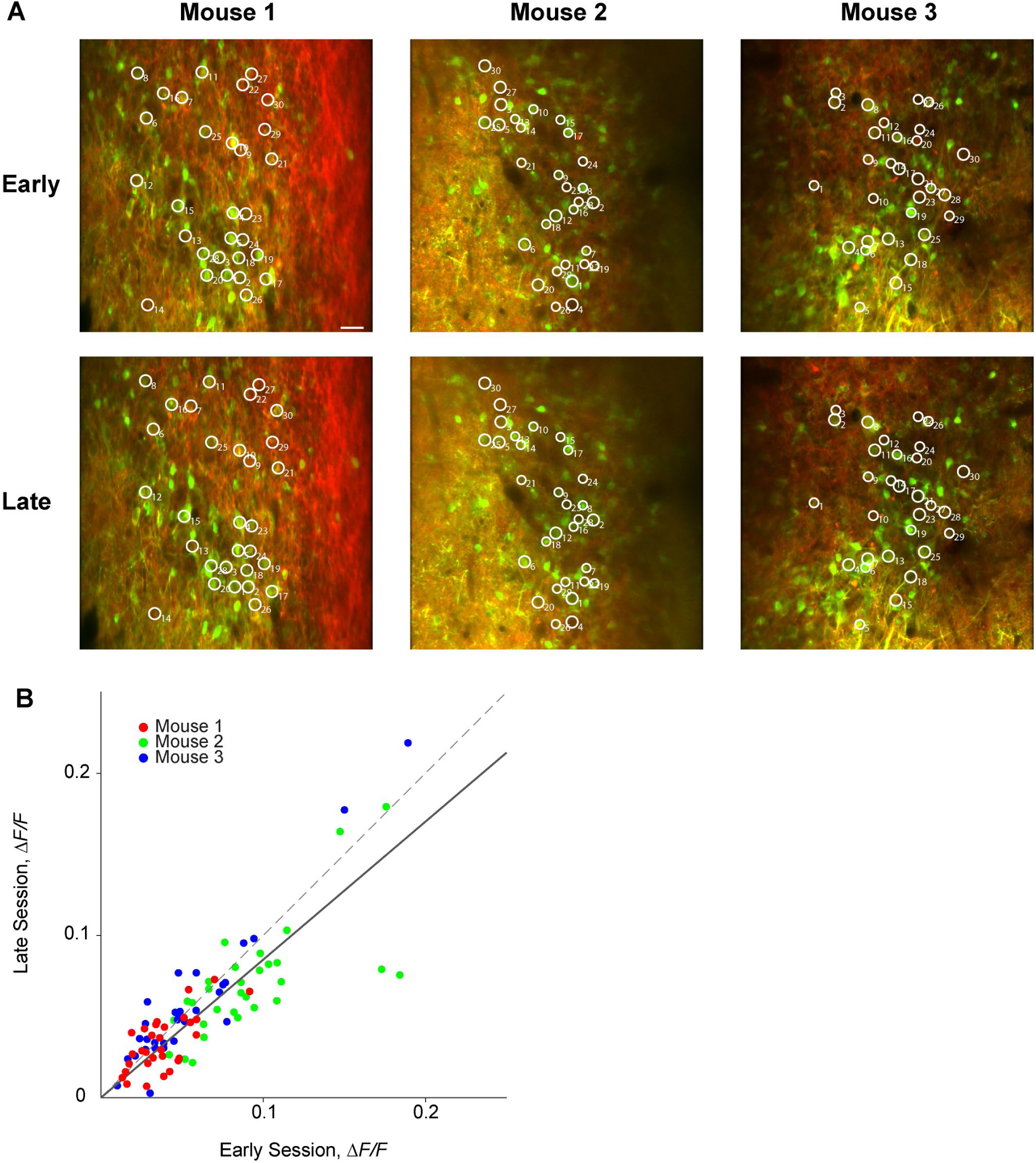
Stability of neuronal targeting and responses across sessions. **A**. Example fields of view and targeted neurons (white circles) imaged early in training (upper) or late in training (lower) for three mice. ChrimsonR-tdTomato expression is shown in red, GCaMP6s expression in green. Scalebar = 30 µm. **B**. Mean response to photostimulation was highly correlated across sessions (30 trials per session, 10 ms stimulation duration, *n*=3 mice, 0.06 mW/µm^2^). Data points represent Δ*F*/*F* values averaged over 330 ms following stimulation. Dashed line is the unity line and the solid line is the linear fit (slope = 0.85).

**Figure S7.**
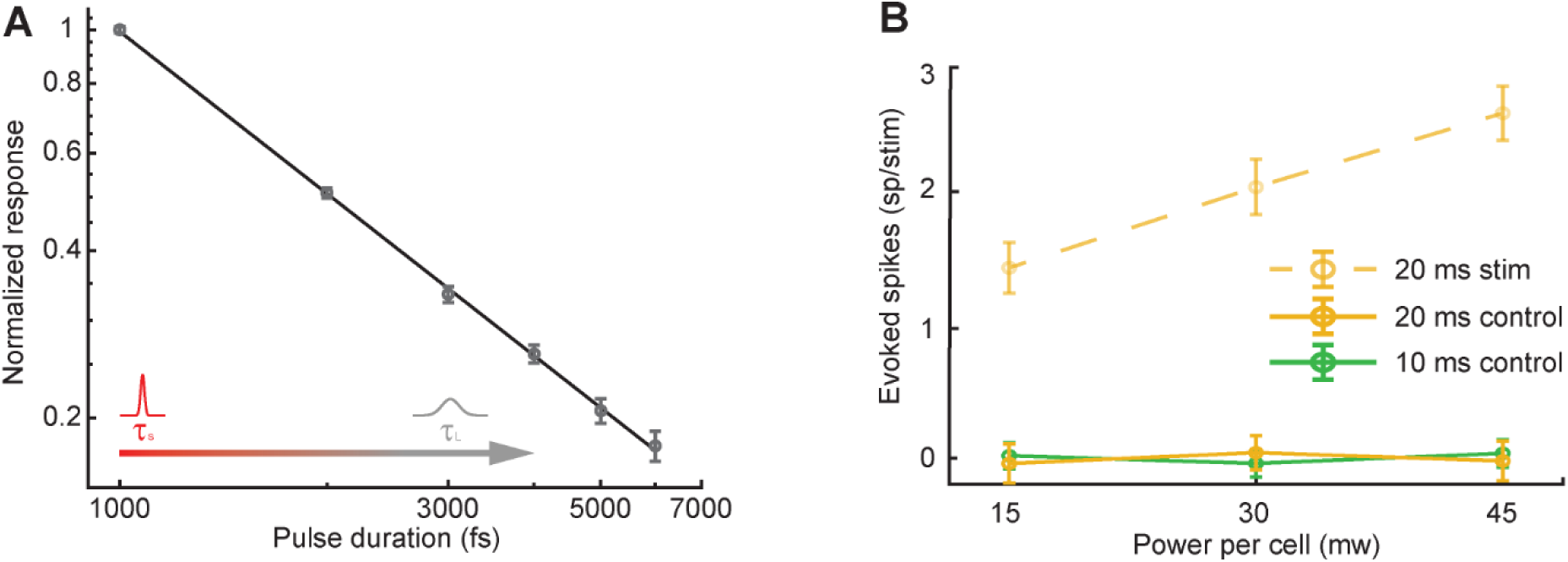
Pulse duration characterization. **A**. Normalized excited 2P fluorescence in fluorescein as a function of pulse width (mean ΔF/F ± s.e.m.). **B.** Average number of spikes evoked by long-pulse sham photostimulation. There was no significant change in the number of evoked spikes compared to baseline (*n* = 4 cells, Wilcoxon rank sum test, *P*=0.99, error bars = ± s.e.m.). The average number of spikes evoked by short pulse photostimulation are shown for comparison (dotted line).

## References

1. Houweling, A.R. and M. Brecht, Behavioural report of single neuron stimulation in somatosensory cortex. Nature, 2008. 451(7174): p. 65-8.

2. Huber, D., et al., Sparse optical microstimulation in barrel cortex drives learned behaviour in freely moving mice. Nature, 2008. 451(7174): p. 61-4.

3. O’Connor, D.H., et al., Neural coding during active somatosensation revealed using illusory touch. Nat Neurosci, 2013. 16(7): p. 958-65.

4. Histed, M.H. and J.H. Maunsell, Cortical neural populations can guide behavior by integrating inputs linearly, independent of synchrony. Proc Natl Acad Sci U S A, 2014. 111(1): p. E178-87.

5. Doron, G., et al., Spiking irregularity and frequency modulate the behavioral report of single-neuron stimulation. Neuron, 2014. 81(3): p. 653-63.

6. Cury, K.M. and N. Uchida, Robust odor coding via inhalation-coupled transient activity in the mammalian olfactory bulb. Neuron, 2010. 68(3): p. 570-85.

7. Shusterman, R., et al., Precise olfactory responses tile the sniff cycle. Nat Neurosci, 2011. 14(8): p. 1039-44.

8. Arneodo, E.M., et al., Stimulus dependent diversity and stereotypy in the output of an olfactory functional unit. Nat Commun, 2018. 9(1): p. 1347.

9. Schneidman, E., et al., Weak pairwise correlations imply strongly correlated network states in a neural population. Nature, 2006. 440(7087): p. 1007-12.

10. Schneidman, E., Towards the design principles of neural population codes. Curr Opin Neurobiol, 2016. 37: p. 133-140.

11. Boyden, E. S., et al., Millisecond-timescale, genetically targeted optical control of neural activity. Nat Neurosci, 2005. 8(9): p. 1263-8.

12. Packer, A.M., et al., Simultaneous all-optical manipulation and recording of neural circuit activity with cellular resolution in vivo. Nat Methods, 2015. 12(2): p. 140-6.

13. Baker, C.A., et al., Cellular resolution circuit mapping with temporal-focused excitation of soma-targeted channelrhodopsin. Elife, 2016. 5.

14. Dal Maschio, M., et al., Linking Neurons to Network Function and Behavior by Two-Photon Holographic Optogenetics and Volumetric Imaging. Neuron, 2017. 94(4): p. 774-789 e5.

15. Shemesh, O.A., et al., Temporally precise single-cell-resolution optogenetics. Nat Neurosci, 2017. 20(12): p. 1796-1806.

16. Yang, W., et al., Simultaneous two-photon imaging and two-photon optogenetics of cortical circuits in three dimensions. Elife, 2018. 7.

17. Forli, A., et al., Two-Photon Bidirectional Control and Imaging of Neuronal Excitability with High Spatial Resolution In Vivo. Cell Rep, 2018. 22(11): p. 3087-3098.

18. Mardinly, A.R., et al., Precise multimodal optical control of neural ensemble activity. Nat Neurosci, 2018. 21(6): p. 881-893.

19. Emiliani, V., et al., All-Optical Interrogation of Neural Circuits. J Neurosci, 2015. 35(41): p. 13917-26.

20. Golan, L., et al., Design and characteristics of holographic neural photo-stimulation systems. J Neural Eng, 2009. 6(6): p. 066004.

21. Rickgauer, J.P. and D.W. Tank, Two-photon excitation of channelrhodopsin-2 at saturation. Proc Natl Acad Sci U S A, 2009. 106(35): p. 15025-30.

22. Andrasfalvy, B.K., et al., Two-photon single-cell optogenetic control of neuronal activity by sculpted light. Proc Natl Acad Sci U S A, 2010. 107(26): p. 11981-6.

23. Papagiakoumou, E., et al., Scanless two-photon excitation of channelrhodopsin-2. Nat Methods, 2010. 7(10): p. 848-54.

24. Dal Maschio, M., et al., Simultaneous two-photon imaging and photo-stimulation with structured light illumination. Opt Express, 2010. 18(18): p. 18720-31.

25. Rickgauer, J.P., K. Deisseroth, and D.W. Tank, Simultaneous cellular-resolution optical perturbation and imaging of place cell firing fields. Nat Neurosci, 2014. 17(12): p. 1816-24.

26. Carrillo-Reid, L., et al., Imprinting and recalling cortical ensembles. Science, 2016. 353(6300): p. 691-4.

27. Paluch-Siegler, S., et al., All-optical bidirectional neural interfacing using hybrid multiphoton holographic optogenetic stimulation. Neurophotonics, 2015. 2(3).

28. Zipfel, W.R., R.M. Williams, and W.W. Webb, Nonlinear magic: multiphoton microscopy in the biosciences. Nat Biotechnol, 2003. 21(11): p. 1369-77.

29. Klapoetke, N.C., et al., Independent optical excitation of distinct neural populations. Nat Methods, 2014. 11(3): p. 338-46.

30. Chen, T.W., et al., Ultrasensitive fluorescent proteins for imaging neuronal activity. Nature, 2013. 499(7458): p. 295-300.

31. Pégard, N.C., et al., Three-dimensional scanless holographic optogenetics with temporal focusing (3D-SHOT). Nature Communications, 2017. 8(1): p. 1228.

32. Osborne, J.E. and J.T. Dudman, RIVETS: a mechanical system for in vivo and in vitro electrophysiology and imaging. PLoS One, 2014. 9(2): p. e89007.

33. Pologruto, T.A., B.L. Sabatini, and K. Svoboda, ScanImage: flexible software for operating laser scanning microscopes. Biomed Eng Online, 2003. 2: p. 13.

34. Gerchberg, R.W. and W.O. Saxton, Practical Algorithm for Determination of Phase from Image and Diffraction Plane Pictures. Optik, 1972. 35(2): p. 237-+.

35. Pnevmatikakis, E.A. and A. Giovannucci, NoRMCorre: An online algorithm for piecewise rigid motion correction of calcium imaging data. Journal of Neuroscience Methods, 2017. 291: p. 83-94.

36. Smear, M., et al., Perception of sniff phase in mouse olfaction. Nature, 2011. 479(7373): p. 397-400.

37. Smear, M., et al., Multiple perceptible signals from a single olfactory glomerulus. Nat Neurosci, 2013. 16(11): p. 1687-91.

